# Allosteric Residue Dynamics in Insulin: Temperature Induces Shift from Dihexamer to Hexamer Collective Motion

**DOI:** 10.1101/2025.01.04.631292

**Authors:** Esra Ayan, Abdullah Kepceoglu

## Abstract

Structures using X-ray diffraction data collected to 2.3 Å, 2.85 Å, and 2.88 Å resolutions have been determined for the long-acting dihexamer insulin at three different temperatures ranging from 100°K to 300°K. It is determined that the unit-cell value of insulin crystal at 100°K temperature has changed at 200°K temperature. This change might be due primarily to subtle repacking of the molecule and loss of noncovalent interactions of myristic acid that binds two hexamers, exhibiting the largest movements. Computational analyses show that allosteric residues and fatty-acid binding residue of insulin hexamers display reduced overall collectivity and inter-residue coupling, likely arising from crystal mosaicity increase and structural fluctuations through elevated thermal motion. This breakpoint has been observed at a characteristic temperature of 200°K, perhaps emphasizing underlying alterations in the dynamic structure of the fatty acid-solvent interface in dimer of hexamer. Combined with computational analysis, findings reveal key insights into thermal stability mechanisms crucial for developing thermostable insulin formulations in industrial applications.

**Teaser:** Temperature-driven shifts in insulin’s structure reveal dynamic changes crucial for enhancing thermal stability in drug formulations.

## Introduction

Our insight into the structural biology of the macromolecular complexes largely relies on X-ray structural studies. The technique provides precise atomic coordinates, and also reveals the temperature factor (Parak et al., 1987). Temperature affects the average structural and dynamic behavior of molecules, making X-ray crystallography an essential tool for generating high-resolution models of molecular structures and analyzing the spatial patterns of atomic motion (Fischer, 2021). By conducting high-resolution X-ray crystallography at varying temperatures, it is possible to investigate molecular surfaces’ physical and chemical features and examine the interplay between bonded and nonbonded forces that maintain structural stability and control molecular dynamics (Massana-Cid et al., 2024). Ancient studies have utilized multi-temperature crystallography to examine how temperature affects the structure and dynamics of various proteins (Frauenfelder et al., 1979; Hartmann et al., 1982; Kuczera et al., 1990). Comparing structures determined through 80K and 300K reveals an anisotropic protein volume expansion of 2-3% with increasing temperature, accompanied by a disorderly rise in atomic motion, as indicated by individual Debye-Waller factors. Studies on this expansion revealed that it primarily involves surface loops, with the additional volume being distributed into subatomic spaces rather than larger, water-sized cavities. The higher Debye-Waller factors observed at 300K were linked to increased vibrational motion and a broader distribution of conformational microstates (Hartmann et al., 1982). Additionally, intermediate temperature structures of a protein have been analyzed (Parak et al., 1982). A study examining the X-ray crystallography of the protein ribonuclease-A across nine different temperatures ranging from 98 to 320 K determined that the protein molecule undergoes a slight expansion with increasing temperature (approximately 0.4% per 100 K), which is linear. At a characteristic temperature around 180–200 °K, the individual atomic Debye-Waller factors exhibit predominantly biphasic behavior, showing a slight positive slope at lower temperatures and a steeper positive slope at higher temperatures (Tilton et al., 1992). Proteins like insulin are highly sensitive to temperature changes (Vimalavathini et al., 2009), and understanding their structural dynamics across a various temperature range is critical for both fundamental and applied research. While the degradation of insulin at elevated temperatures is well-documented (M. Rasmussen et al., 2024), the specific structural changes and dynamics associated with varying temperatures, particularly in long-acting formulations, remain less explored. For this study, the structural biology of the detemir insulin has been investigated since detemir insulin comprises two adjacent hexamers by conjugating myristic acid; thus, each hexamer has primarily a-helices packed around two additional zincs and has no cofactor (Home et al., 2006). The structural and ligand-binding characteristics of hexamer insulin have been extensively studied (Hassiepen et al., 1999). Insulin molecules assemble into dimers in the presence of zinc ions, further organized into hexamers composed of three insulin dimers arranged symmetrically around two zinc ions along a 3-fold axis (Brange et al., 1993). Each zinc ion coordinates with three B10 histidine imidazole groups, and additional extrinsic ligands complete the coordination, resulting in either tetrahedral or octahedral ligand geometry. The three-dimensional structures of insulin in its monomeric, dimeric (Ciszak et al., 1994; Gursky et al., 1992), hexameric (Smith et al., 2000), as well as dihexameric (Whittingham et al., 1997) forms have been characterized using X-ray crystallography. Hexamer insulin comprises six homo-monomer subunits that each monomer includes two polypeptide chains (the A-chain and the B-chain). The A-chain, which includes 21 amino acids, features two short α-helices, spanning residues A1–A8 and A13–A19. The B-chain, containing 30 amino acids, centers around a primary helix (B9–B19) with regions of extended structure at each end. Two interchain disulfide bonds, A7–B7 and A20–B19, securely connect the chains, while an additional intrachain disulfide bond links A6 to A11 within the A-chain (Ayan & DeMirci, 2022). Hexamer insulin can be engineered to synchronize its release from the injection site with the body’s physiological requirements, allowing control over both release and absorption rates (Rosskamp et al., 1999) Beyond the fast-acting hexamer form, a more prolonged action profile for dihexamer insulin can be achieved by incorporating albumin-binding groups, which help extend its circulation time in the bloodstream (Home et al., 2006). Research interest in insulin is driven by its pharmaceutical importance and application as a model for understanding allosteric protein behavior (Jarosinski et al., 2021). This study aims to investigate the effect of temperature on the allosteric sites of dihexamer/hexamer insulin and if temperature changes influence the crystal lattice and mosaicity of the dihexamer insulin crystal. X-ray crystallography was performed on a cryogenic single insulin crystal (Fig. S1), with data collected at 100°K, 200°K, and 300°K to introduce temperature jumps, enabling a comprehensive computational analysis of structural dynamics at those temperatures. Temperature-jump single-crystal X-ray crystallography offers valuable insights into the quaternary structure of dihexamer/hexamer insulin, specifically highlighting the allosteric residue dynamics and temperature-dependent behaviors characteristic of its oligomeric form.

## Results and discussion

### Temperature shift alters the unit cell of insulin crystal

This session of the study yielded comprehensive insights into temperature-dependent structural transformations in the dihexameric insulin crystal (Table 1). Data collected from a single cryogenic crystal (Fig. S1) at incremental temperatures (100 °K, 200 °K, and 300 °K) revealed significant shift the unit cell parameters from dihexamer to hexamer forms (Fig 1). The insulin crystal structure at 100 °K, 200 °K, and 300 °K respectively illustrates significant changes in unit cell dimensions as the temperature increases. At 100 °K, the unit cell exhibits idealized rhombohedral symmetry with parameters a = b = c = 79.0 Å, = = 90°, = 120°, indicative of a highly ordered lattice configuration (Fig. 1A). This cryogenic condition preserves the integrity of the original rhombohedral unit cell, minimizing thermal perturbations (Kurinov et al., 1995). However, at 200 °K, there is a notable transformation, as the unit cell shifts to a = b = 80.7 Å, c = 39.6 Å, = = 90°, = 120° (Fig. 1D). This change, particularly the reduction in the c-axis, suggests a structural rearrangement due to the increased thermal motion, affecting intermolecular distances along this axis and indicating some type of phase transition (Parak et al., 1987). At 300 °K, the unit cell further contracts to a = b = 78.3 Å, c = 39.6 Å, = = 90°, = 120°, indicating sustained thermal effects along the a and b axes and stabilization of the c-axis contraction (Fig. 1G). This anisotropic contraction suggests that temperature fluctuations alter the native hexameric rhombohedral structure, creating temperature-induced strain within the lattice that may influence molecular packing and overall lattice energy (Cucka et al., 1970; Juers et al., 2001; Kihara, 1978; Stevens et al., 2015; Tilton et al., 1992). The electron density maps at 100 °K, 200 °K, and 300 °K temperatures underscore the effect of thermal energy on the spatial distribution of electron density. At 100 °K (Fig. 1B), the electron density is sharply defined at a resolution of 2.3 Å, with well-resolved contours indicating minimal atomic displacement. This observation closely aligns with thermal displacement analyses discussed later (Fig. 2). The crisp electron density highlights a densely packed lattice with strong intermolecular interactions (Novikov et al., 2020; Saldivar-Garcia et al., 2004), characteristic of cryogenically stabilized systems (Novikov et al., 2020). Upon increasing the temperature to 200 °K at 2.83 Å resolution (Fig. 1E), the electron density contours begin to exhibit mild blurring in specific regions, suggesting an incremental increase in atomic displacement and dynamic disorder. This thermal relaxation effect is especially prominent along the c-axis, where the unit cell contraction may reduce intermolecular contact and loosen molecular packing (Juers et al., 2001; Tilton et al., 1992). At 300 °K (Fig. 1H), the electron density map displays significant diffusion, with widespread contour blurring across the structure, indicating enhanced atomic mobility (Fig. 2E). The pronounced electron density distortion at higher temperatures reflects a marked increase in intramolecular and intermolecular flexibility (Fig. S2), signaling that thermal agitation at 300 °K disrupts the lattice coherence, likely leading to local conformational changes and decreased structural rigidity (Tilton et al., 1992; Young et al., 1994). Additionally, at 100 °K, an RMSD of 0.298 reflects high structural stability (Fig. 1C). With increasing temperature, the RMSD rises to 0.445 at 200 °K and 0.517 at 300 °K (Fig 1F-I), supporting progressively greater thermal instability (Fig. 2), interchain misalignment, and enhanced flexibility and mobility at allosteric sites.

**Figure 1.**
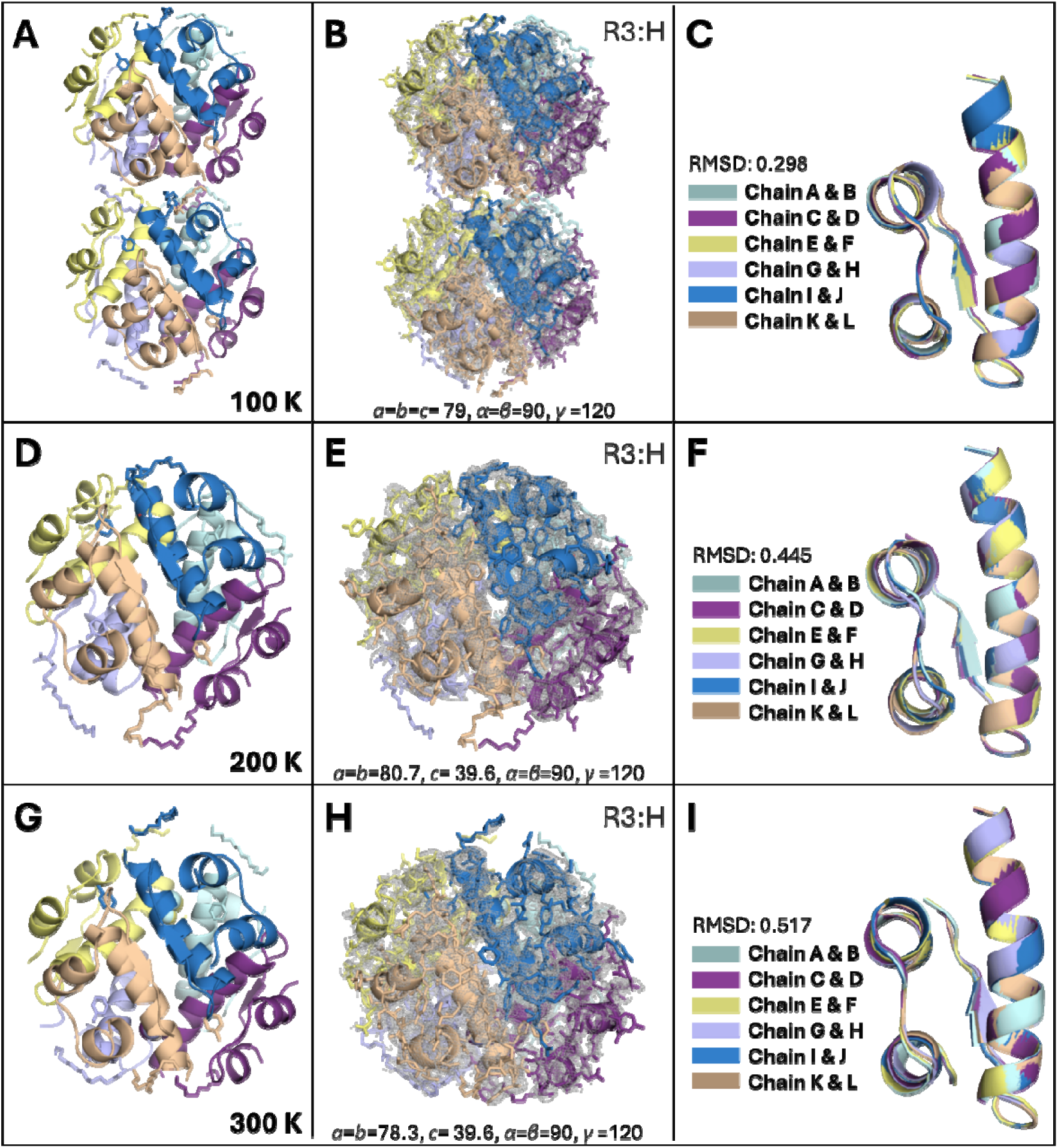
Structural comparison of dihexamer and hexamer insulin crystals at 100 K, 200K and 300K. (A, D, G) Structures of dihexamer (A) and hexamer (D, G) insulin are colored based on each chain of hexamer. **(B, E, H)** 2*F*o-*F*c simulated annealing-omit map at 1 sigma level is colored in gray. The unit cell =90, β =120 for dihexamer at 100 °K (E) a=b=80.7, c= 39.6, α=β γD=120 for hexamer at 200 °K (H) a=b=78.3, c= 39.6, α=β=90, γD=120 for hexamer at 300 °K **(C, F, I)** Each chain of insulin structures at 100°K, 200 °K and 300 °K is superposed with an overall RMSD of 0.298DÅ, 0.445 Å, and 0.517 Å, respectively.

**Figure 2.**
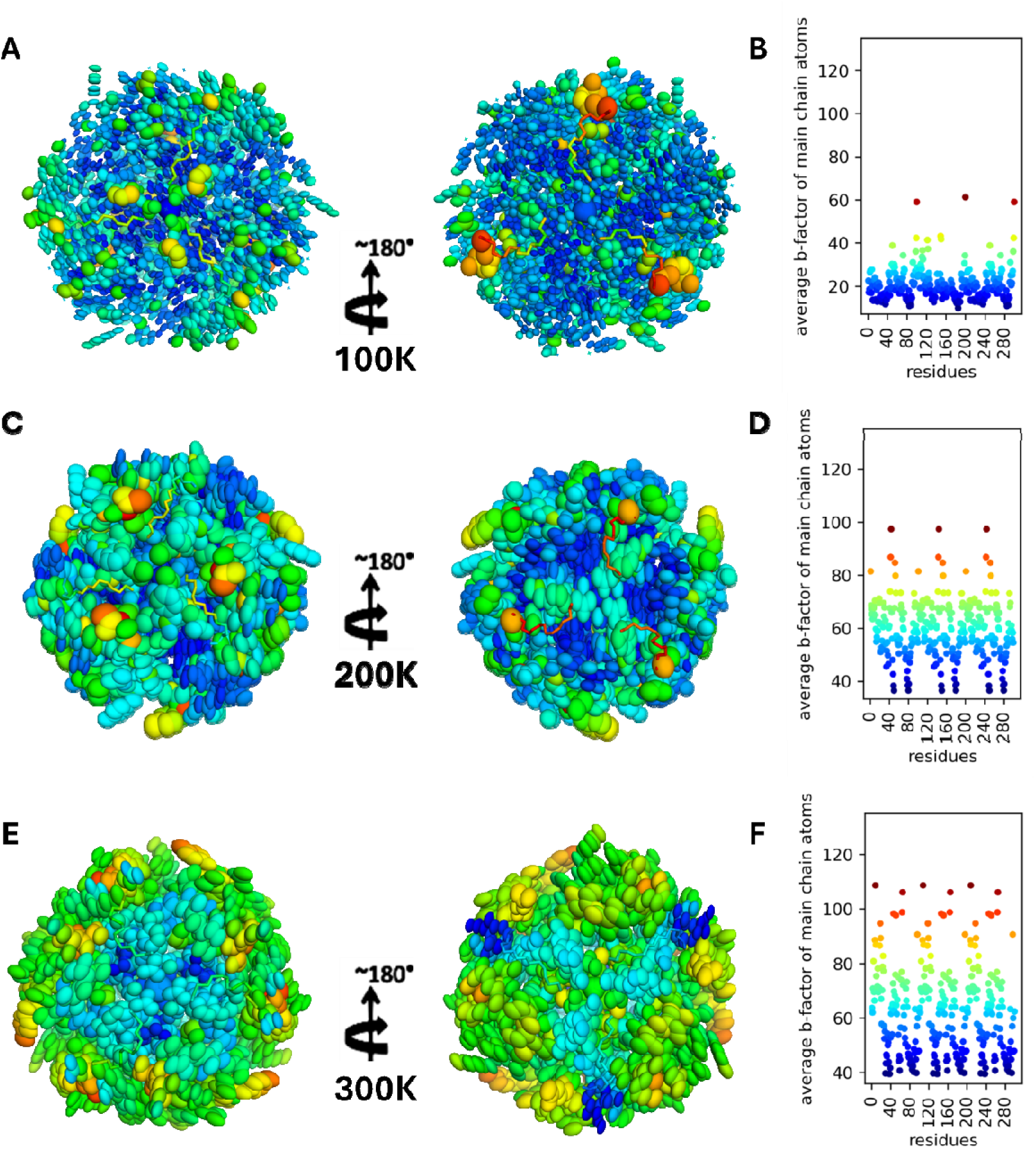
Comparative ellipsoids representation and *B*-factor (Å^2^) plots of hexamer portion of insulin structures at various temperatures. (**A**) Thermal displacement analysis by ellipsoids representation, and *B*-factor plot **(B)** of hexamer insulin at 100 **°**K temperature. **(C)** Thermal displacement analysis by ellipsoids representation, and *B*-factor plot **(D)** of hexamer insulin at 200 **°**K temperature. **(E)** Thermal displacement analysis by ellipsoids representation, and *B*-factor plot **(F)** of hexamer insulin at 300 **°**K temperature. On the y-axis are plotted the *B*-factors of the main chain atom, averaged per residue.

**Table 1.** Data collection and refinement statistics.

| **Data collection** | **Dihexamer – 100K** | **Hexamer – 200K** | **Hexamer – 300K** |
| --- | --- | --- | --- |
| **Instrument** | Turkish Light Source | Turkish Light Source | Turkish Light Source |
| **Resolution range** | 19.77- 2.3 (2.382- 2.3) | 17.56 - 2.85 (2.951 - 2.85) | 17.11- 2.881 (2.983 - 2.881) |
| **Space group** | R 3: H | R 3: H | R 3: H |
| **Unit cell** | 79.06 79.06 79.49 90 90 120 | 81.23 81.23 40.32 90 90 120 | 78.31 78.31 39.63 90 90 120 |
| **Redundancy** | 4.1 (4.0) | 3.4 (3.0) | 2.9 (1.9) |
| **Completeness (%)** | 98.77 (98.67) | 98.53 (94.01) | 83.79 (86.36) |
| **Mean I/sigma(I)** | 30.01 (14.02) | 6.46 (2.35) | 3.6 (1.53) |
| **Wilson B-factor** | 16.65 | 57.39 | 51.00 |
| **R-merge** | 0.03625 (0.08822) | 0.06099 (0.099) | 0.170 (0.022) |
| **R-meas** | 0.0417 (0.1021) | 0.06244 (0.0934) | 0.319 (0.024) |
| **R-pim** | 0.0203 (0.05086) | 0.0594 (0.068) | 0.0977 (0.012) |
| **CC1/2** | 0.999 (0.99) | 0.934 (0.99) | 0.947 (0.999) |
| **CC\*** | 1 (0.998) | 0.983 (0.704) | 0.975 (0.609) |
| **R-work** | 0.2336 (0.2560) | 0.2854 (0.3990) | 0.3189 (0.4410) |
| **R-free** | 0.2881 (0.2676) | 0.3369 (0.4509) | 0.3912 (0.6081) |
| **No. of reflections** | 33832 (3265) | 48296 (3457) | 36211 (2926) |
| **Unique reflection** | 8201 (813) | 2306 (202) | 1729 (171) |
| **Protein residues** | 200 | 100 | 100 |
| **RMS (bonds)** | 0.002 | 0.002 | 0.002 |
| **RMS (angles)** | 0.38 | 0.35 | 0.39 |
| **Ramachandran favored (%)** | 100.00 | 100.00 | 100.00 |
| **Ramachandran allowed (%)** | 0.00 | 0.00 | 0.00 |
| **Ramachandran outliers (%)** | 0.00 | 0.00 | 0.00 |
| **Average B-factor** | 24.37 | 58.58 | 72.49 |
| **macromolecules** | 23.13 | 58.81 | 72.67 |
| **ligands** | 45.32 | 54.62 | 71.66 |

Thermal displacement (*B*-factor) analysis by representing ellipsoid mode of insulin dihexamer/hexamer was performed at three distinct temperatures (Fig. 2). The experimental investigation of dynamic equilibrium fluctuations in proteins is made possible through the measurement of the Debye-Waller or B-factor for individual atoms using X-ray crystallography (Kuzmanic et al., 2010). Such analysis can highlight the dynamic behavior and residue mobility of the protein under varying thermal conditions. At 100 **°**K, the thermal displacement parameters (*B*-factors) are minimal, indicating that the) atoms are well-ordered with restricted thermal motion. (Fig. 2A) The ellipsoid representation in Panel A shows a dominantly blue-green narrow ellips, suggesting lower mobility across large variety of residues. The graph in Panel B is consistent with this ellipsoid representation even, with excess residues indicating lower *B*-factors largely clustered within the lower range of the *B*-factor spectrum (1.00 Å^2^ – 60.00 Å^2^). Only a few residues display moderate *B*-factor values, represented as yellow-orange-red regions, suggesting localized mobility at those residues (T27_B_-P28_B_-K29_B_) act as a molecular switch, facilitating the transition from the dihexameric to the hexameric form. At 200 **°**K, there is an observable increase in thermal motion, as evidenced by the ellipsoid representation in Fig. 2C, where most regions exhibit green and yellow relatively vibrate spheres, indicating increased mobility compared to 100 **°**K (Fig. 2A), but not as much as at 300 **°**K (Fig. 2E). The corresponding graph in Fig. 2D shows a broader distribution of *B*-factors than 100 **°**K, with several residues experiencing moderate-to-high flexibility. This temperature introduces apparent dynamic effects, particularly in specific regions (G1_A_, I2_A_, V3_A_, R22_B_, and G23_B_), possibly correlating with the onset of hexamerization and disruption of dihexamer stability. This is compatible with the literature claiming the temperature of 180-220 **°**K is the breaking point of fundamental changes in the dynamic structure of the surrounding protein solvent (B. F. Rasmussen et al., 1992; Tilton et al., 1992), which can also explain why unit cell changing is observed at 200 **°**K temperature. At 300 **°**K, thermal displacement parameters are significantly higher, indicating improved atomic motion and greater residue mobility (Fig. 3E-F). The ellipsoids representation in Panel E shows substantial yellow and orange vibration, with distinct areas displaying red, signifying higher residue mobility. The graph in Panel F reveals a wide range of *B*-factors, with substantial residues exceeding the threshold of coherent residue blocks. This behavior aligns with the transition to a hexameric form, where increased temperature destabilizes the dihexameric quaternary interactions (Kontopoulos et al., 2020; Zheng et al., 2024), enhancing residue-level dynamics and reduced collective motion (Medeiros Almeida et al., 2021).

**Figure 3.**
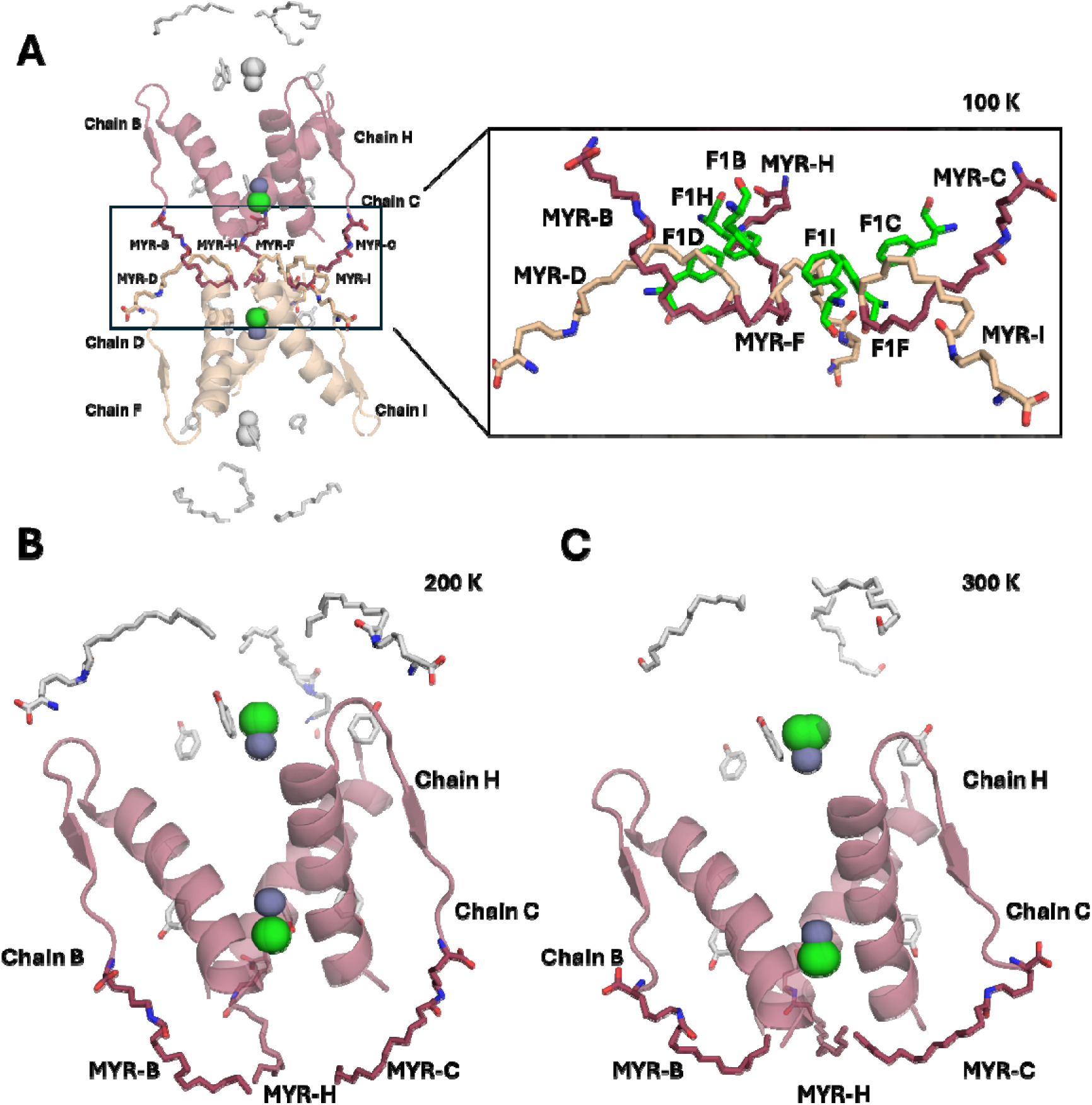
Cartoon representation and localization of MYR moiety on dihexamer/hexamer insulin structures at different temperatures. **(A)** MYR localization of dihexamer insulin structure at 100 °K. Two hexamers were integrated by MYR chains to form dihexamerization; MYR moieties that facilitate integration of one hexamer (Chains B, C, and H) are colored raspberry and others in the other hexamer (Chain D, F, and I) are colored wheat. MYR molecules that facilitate dihexamerization are shown in close-up view in the right panel. Phe 1 residues that facilitate to establish hydrophobic interaction with MYR are colored in green. The 200 °K **(B)** and 300 °K **(C)** structures are shown in panel B and panel C, respectively. MYR molecules that facilitate dihexamerization are colored raspberry.

### Myristic acid affects temperature-dependent crystal packing dynamics

To analyze the impact of increasing temperature on the transition from dihexamer to hexamer, focused on the role of myristic acids based on insulin structure and thermodynamic implications (Fig. 3 and Fig. S2). The dihexameric insulin structure is stabilized at cryogenic temperature (Fig. 3A) by hydrophobic solid interactions and a well-organized arrangement of myristic acid (MYR) moieties, indicating that MYR stabilizes the dihexameric structure by maintaining both hexamers’ integrity (Bech et al., 2018). This is in line with the electron density for MYR molecules, which is well-defined, with continuous density along the entire length of the fatty acid chains (Fig. S2A). At 200 °K, discern that shifting the dihexamer detemir to its hexamer form (Fig. 3B) is probably due to loss of stability as the elevated temperature can disrupt the hydrophobic interactions (Sun et al., 2022; Wang et al., 2014) between myristic acid and the side chain of FB1 (Phe in chain B) (Whittingham et al., 1997) facilitated by the crystal mosaicity (see, Table 1 “Average B-factor” toward higher temperature) (Lakhloufi et al., 2018) and lattice expansion that disrupts intermolecular interactions (Frauenfelder et al., 1987; Saldivar-Garcia et al., 2004; Zhuravleva et al., 2015). It is consistent with the electron density maps revealing largely disorder in the myristic acids (Fig. S2B), suggesting that dissociating the dihexameric insulin to hexamer form might begin with the Brownian motions and fluctuations of the myristic acid (Ayan et al., 2023) due to loss of water molecules at elevated temperature. It is also consistent with thermal displacement analysis (Fig. 2A-B), which shows that P28_B_-K29_B_ –T30_B_ residues, essential residues for the initial dissociation process (Holleman, 2015; Samuel et al., 2008; Whittingham et al., 1997), display higher fluctuation at cryogenic temperature even. At 300 °K, the increased thermal motion stabilizes this significant conformational shift (Fig. 3C). Integrating structural and electron density data can propose clear evidence for the temperature-dependent dynamics of myristic acids and their pivotal role in the dihexamer-to-hexamer transition.

### Computational analyses confirm temperature-dependent dynamics of insulin

Protein binding occurs by forming new non-covalent interactions between specific residues of two proteins. These interactions can alter the dynamics of residues at distant sites by disrupting existing communication pathways and establishing new ones. Such rearrangements have the potential to significantly reshape the traditional residue communication network. Normal Mode Analysis (NMA) can effectively detect and analyze these changes (Eren et al., 2021). GNM (Gaussian Network Model) (Bahar et al., 1997)and ANM (Anisotropic Network Model) (Atilgan et al., 2001) based on NMA are particularly effective in predicting large-scale protein motions in slow modes and localized dynamics in fast modes (Bahar et al., 1998). In slow modes, regions with higher fluctuations usually indicate flexible, mobile segments, while regions with minimal fluctuations correspond to hinge points critical for protein function (Yang et al., 2005). Otherwise, fast modes highlight residues with peak fluctuations, often identifying structurally constrained hotspots required for processes such as binding or folding (Demirel et al., 1998). Moreover, analyzing changes in residue dynamics in all modes provides important information about proteins’ functional mechanisms and allosteric regulation (Ayan, Yuksel, et al., 2022). Accordingly, extensive GNM analysis was performed to analyze large-scale oligomeric insulin motions in slow modes and localized dynamics in fast modes, toward to higher temperatures (Fig. 4). The temperature-dependent collective motions analysis of the insulin crystal data was performed to reveal if the presence of significant alterations in the residue dynamics of insulin structures through different oligomeric states and temperatures (Fig. 4A). At 100°K, the reference dihexamer model (PDB ID: 1XDA) shows high collectivity for certain slow modes (close to 1.00 au). Comparatively, collected dihexamer at 100°K shows a similar trend but highlights slight variances, particularly in the distribution of collectivity, suggesting minor deviations likely arising from lower crystal mosaicity increase (Lakhloufi et al., 2018; Meents et al., 2010) and structural fluctuations. Toward higher temperature values, the transition to the hexameric form is in line with apparent shifts in collective motions. At 200°K, the hexamer data displays enhanced collectivity in certain modes while reducing cooperation in others (close to 0.7 au), suggesting a rearrangement of the residue dynamics within the hexameric assembly. This shift emphasizes the temperature-dependent behavior of oligomeric insulin, where specific residues display greater adaptability at intermediate thermal conditions (Somero, 2003; *Thermostability of human insulin Background*, n.d.). At 300°K, the hexamer keeping oligomeric, but exhibits reduced overall collectivity compared to the 200°K one. This shift in collectivity suggests diminished cooperative dynamics, likely due to increased thermal motion and reduced inter-residue coupling. These findings collectively highlight the temperature-sensitive allosteric topography of insulin, with structural dynamics tied to oligomeric state and thermal conditions (Baheri et al., 2016).

**Figure 4.**
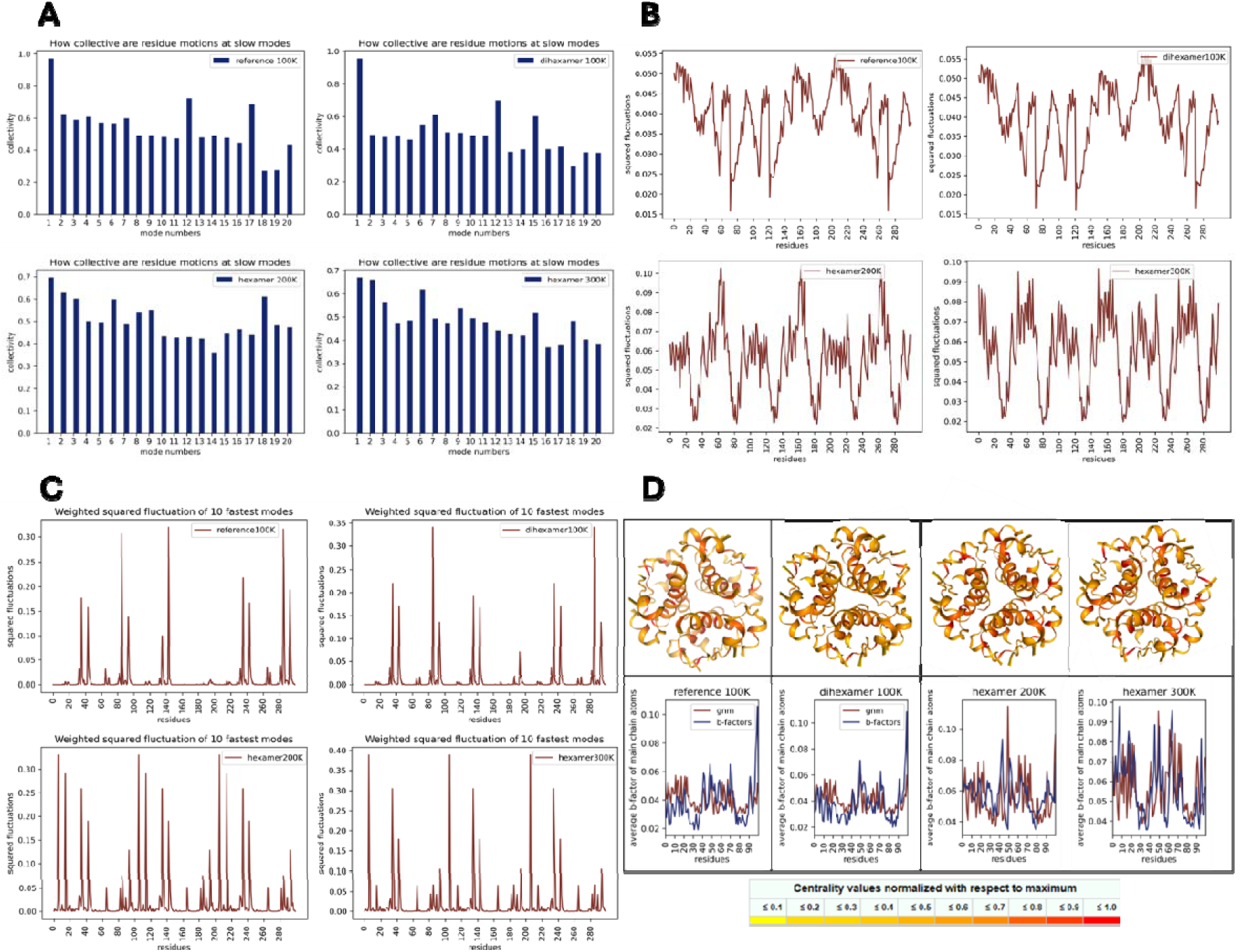
Normal mode analysis and network-based analysis of dihexamer (at 100°K) and hexamer insulin structures (at 200°K then 300°K). **(A)** Slow mode-based collectivity analysis of reference model (PDB ID: 1XDA), dihexamer at 100°K, and hexamer structures at 200°K then 300°K temperatures, respectively. **(B)** Comparisons the weighted squared fluctuation of ten slowest modes for reference model (PDB ID: 1XDA), dihexamer at 100°K, and hexamer structures at 200°K then 300°K temperatures, respectively. **(C)** Comparisons the weighted squared fluctuation of ten fastest modes for reference model (PDB ID: 1XDA), dihexamer at 100°K, and hexamer structures at 200°K then 300°K temperatures, respectively. **(D)** Node centrality results from NAPS and computational/experimental b-factor results from GNM of oligomeric insulin complexes in different temperatures. Hexamer insulin structures in cartoon representation are colored by degree centrality. In *B*-factor plot, computational one is colored in raspberry and experimental one is colored in sky blue.

The weighted squared fluctuation of the ten slowest modes from the insulin crystal structures at different temperatures highlights the effect of thermal changes on residue dynamics in the slow modes (Fig. 4B). The reference dihexamer model and our cryogenic dihexamer data at 100°K exhibit fluctuations characteristic of the stable, cryogenic oligomeric state. Regions with diminished higher fluctuations (maxima loci) align with mobile loop regions or solvent-exposed residues, while the largely observed lower fluctuations (minima loci) mark dynamic dissections critical for the receptor binding sites and allosteric function of the insulin (Eren et al., 2021). At 200°K, the hexameric form shows an obvious increase compared to cryogenic structures in fluctuation through many regions, which is in line with the loop regions or solvent-exposed residues. This temperature-induced transition dihexamer to the hexamer form likely had enhanced the mobility of myristic acid (Bedford et al., 123 C.E.; Sarı et al., 2001) and its binding residue, which might indicate a partial destabilization of the oligomeric assembly. The hinge points critical for structural stability, especially myristic acid binding residues on slow mode plot, show fewer fluctuation increases, keeping their functional roles despite the elevated temperature. The 300°K hexamer data, however, reveals a further increase in fluctuation, particularly in some hinge segments at 200°K which is also characterized by receptor interaction sites of insulin. The mean square displacements of individual residues through 300°K to 100°K exhibit an apparent linear decline as temperature decreases, suggesting that the potential governing their motions follows a parabolic profile (Parak et al., 1987).

Also, the weighted squared fluctuation of the ten fastest modes was calculated using those structures (Fig. 4C). The analysis of fast modes reveals significant temperature-dependent dynamics in residue fluctuations between cryogenic dihexamer and moderate then ambient hexamer structures, highlighting structurally critical hotspots for binding, folding, and allosteric regulation processes (Eren et al., 2021). At 100°K, the reference dihexamer model exhibits limited fluctuation peaks, aligning with rigidly restricted regions essential for keeping structural stability under cryogenic conditions. The experimental 100°K dihexamer data closely mimics the reference, with minor variations in peak amplitudes and positions around the 130-150 residues, likely due to intrinsic crystal mosaicity (Meents et al., 2010), while preserving key hotspot regions. At 200°K and 300°K, shifting the hexameric form, increased fluctuation amplitudes, and enriched novel peak oscillations have been observed in the fast modes. Notice that around the residues 120 and between residues 200 and 240 in the cryogenic dihexamer forms have experienced a further reduction than their hexamer forms, suggesting the residues in hexamer forms gain their kinetic hotspot status while restricted in dihexamer forms. The enriched hotspots that emerge in the hexamer structure may be interpreted as the favorable interaction within oligomer insulin that has been initialized by shifting the dihexamerization-to-hexamerization process. Peak dynamics profile between the 200°K and 300°K hexamer structures supported the further elevated amplitudes (close to 0.4 au) that emerge in the hexamer structure at 300°K, consistent with significant kinetic energy variabilities observed at elevated temperatures (Tilton et al., 1992). Findings collectively highlight more plausible dynamic adaptability of insulin hexamer at higher temperatures than under cryogenic conditions, as well as the redistribution of hotspots characterized by the functional significances of temperature-induced mobility was also supported by enhanced kinetic variability of the hexamer at 300°K temperature than 200°K temperature one (Fig. 4C).

Proteins have traditionally been analyzed based on their secondary structure and fold architecture; however, representing proteins as networks of non-covalent interactions between amino acid residues offers a systems-level approach to understanding the topological features of complex three-dimensional structures, where the centrality of a residue reflects its structural and functional significance within the residue-residue contact network (Bagler et al., 2007; Blacklock et al., 2014; Brinda et al., 2005; Taylor, 2013). The network-based analysis of protein structures (NAPS) was performed to understand the quantitative/qualitative mechanism of residue-residue interactions in oligomeric insulin structures (Chakrabarty et al., 2016). Figure 4, panel D and Table 2 summarize the structural conformation of insulin from dihexamer to hexamer forms through varying temperatures, focusing on degree analysis of the node centrality and global parameters from NAPS. Dihexamer structure at 100°K shows the highest node number with a degree centrality (27 nodes >11), indicating a more collective network of highly connected residues than the reference dihexamer (21 nodes). Temperature-induced shifting to hexamer at 200°K, notable reduction is observed in the number of nodes exceeding the thresholds for closeness (9 nodes >0.18) and betweenness (21 nodes >0.4), suggesting relatively less collective network with shorter paths (average shortest path = 5.94). This is in line with the reduction in diameter (11.00), reflecting a decreased structural reorganization and residue interactions (Chakrabarty et al., 2016). Otherwise, network of the hexamer at 300°K becomes more adaptive, supported by fewer nodes meeting the centrality thresholds. The number of nodes with a degree >11 decreases to 18, while nodes exceeding closeness >0.18 drop significantly to 3, highlighting increased mobility and reduced centralized control. All analyses from GNM and NAPS suggest shifting from dihexamer to hexamer, coupled with increasing temperature, results in intrinsic dynamics that expand the protein molecule, reflecting structural adaptations that optimize connectivity and mobility in response to thermal motion (Frauenfelder et al., 1987; Kurinov et al., 1995; Parak et al., 1987; Tilton et al., 1992; Young et al., 1994).

**Table 2.** Centrality parameters provided in NAPS for dihexamer and hexamer structures.

| Global parameters | Structures |  |  |  |
| --- | --- | --- | --- | --- |
|  | PDB ID: 1XDA | Dihexamer 100°K | Hexamer 200°K | Hexamer 300°K |
| **Nodes:** | 300 | 300 | 300 | 300 |
| **Edges:** | 1146 | 1158 | 1164 | 1143 |
| **Diameter** | 12.00 | 13.00 | 11.00 | 12.00 |
| **Radius** | 9.00 | 9.00 | 9.00 | 9.00 |
| **Avg degree** | 7.64 | 7.72 | 7.76 | 7.62 |
| **Avg shrt paths** | 6.40 | 6.42 | 5.94 | 6.05 |
| **Clst coeff.** | 0.53 | 0.53 | 0.53 | 0.54 |
| **Node centrality > Degree** | 21 nodes (>11) | 27 nodes (>11) | 21 nodes (>11) | 18 nodes (>11) |
| **Node centrality > Closeness** | 18 nodes (>0.18) | 15 nodes (>0.18) | 9 nodes (>0.18) | 3 nodes (>0.18) |
| **Node centrality > Betweenness** | 21 nodes (>0.4) | 21 nodes (>0.4) | 21 nodes (>0.4) | 17 nodes (>0.4) |
| **Eigenvectos centrality** | 18 nodes (>0.79) | 21 nodes (>0.79) | 18 nodes (>0.79) | 12 nodes (>0.79) |

To illuminate if temperature changes affect interface interaction between monomers of dihexamer and hexamer structures, cross-correlation analysis from GNM was performed. GNM also captures the collective motions of residues by the top slowest modes together with cross-correlation, enabling a more detailed explanation of the changes that occur at both interface and distal sites during binding. Cross-correlation heatmap at 100°K, 200°K, and 300°K reveal significant shifts in residue coupling as the temperature increases (Fig. 5). At 100°K, intermonomer residues (between residues 200 and 300) exhibit high collectivity (cross-correlations close to 1 (red)), reflecting more coherent movements, means that dihexamer structure imports a higher stabilization (Parak et al., 1987). Slow modes along the same residue regions further identify fewer dynamic dissections on cryogenic oligomer, with the minima in the slow mode trend highlighting critical residues for protein function. Both cross-correlation values from GNM at 200°K and 300°K indicate more deviations in the directions of distinct residue motions moving in opposite directions isolated by several hinges. This is in line with slow mode analysis from GNM and consistent with thermal motion disrupting the collective behavior at the distal sites and interface relations. Obviously, collective regions progressively lose their cohesive dynamics, thus stressing less intermonomer relations toward elevated temperature (B. F. Rasmussen et al., 1992; Tilton et al., 1992). These findings reveal insulin hexamer undergoes temperature-dependent dynamic variations in intermonomer collectivity, shifting from coherent movements at cryogenic temperatures to mobile and less cohesive interrelations toward elevated thermal motion.

**Figure 5.**
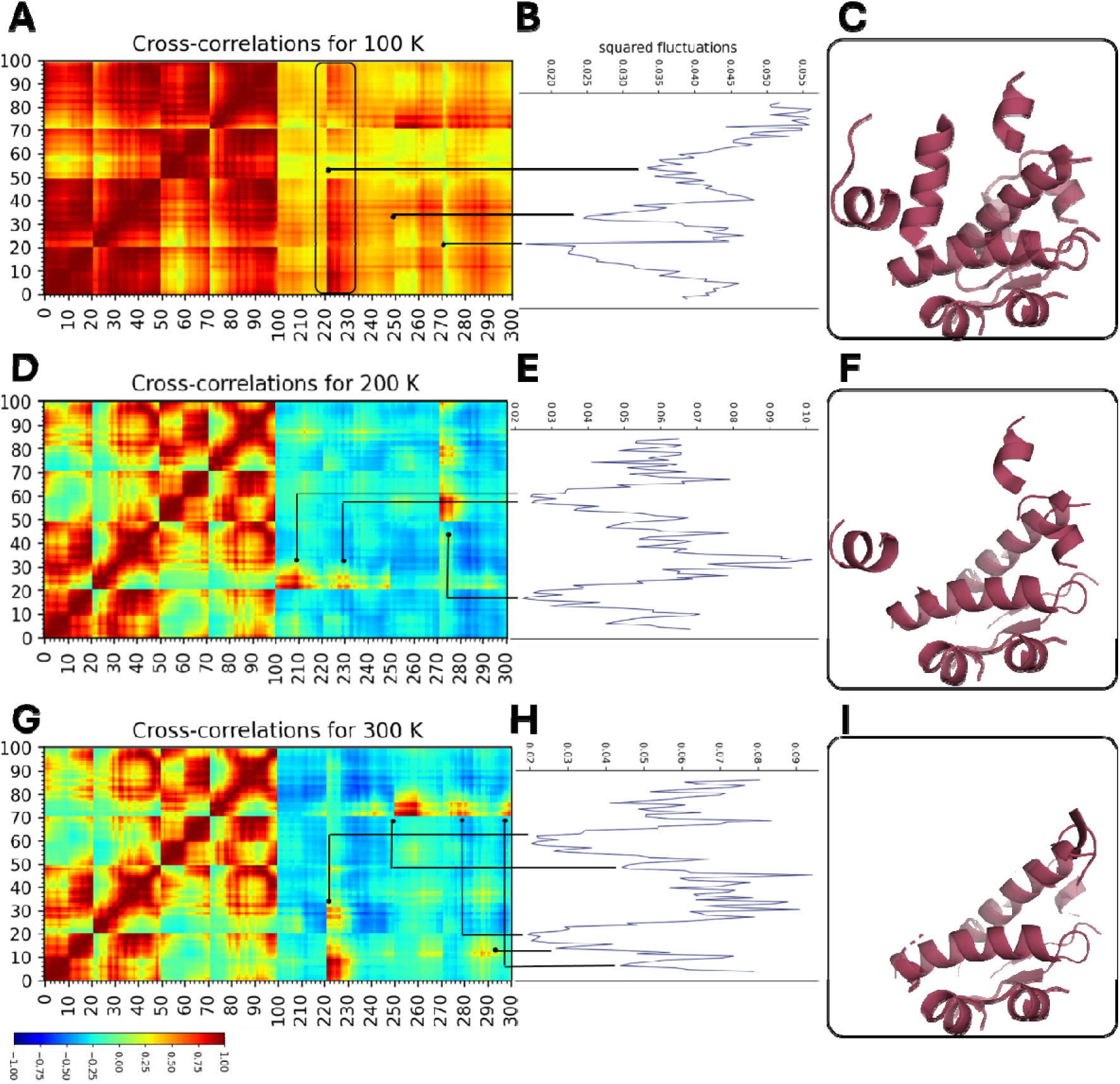
Comparison of cross-correlation map and slow modes from GNM of dihexamer and hexamer structures at various temperatures. **(A)** Cross-correlation for dihexamer structure at 100°K shows strong collectivity at the interface. This is supported by **(B)** slow modes analysis of the interface region. Potential hinges have been marked by arrows. Interface regions of cryogenic large oligomer have a strong correlation, which is visualized by **(C)** cartoon representation. **(D)** Cross-correlation for hexamer structure at 200°K is less coherent at the interface than cryogenic oligomer. Especially, interface sites display weak correlations and are separated by more hinges in slow modes **(E).** Obvious hinges have been marked by arrows. Interface regions of 200K oligomer have a weak correlation, which is visualized by **(F)** cartoon representation. **(G)** Cross-correlation for hexamer structure at 300°K is less coherent and shows more inverse correlations isolated by hinges at the interface. **(H)** This is in line with the slow modes trend. Obvious hinges have been marked by arrows. Interface regions of hexamer at ambient temperature have very weak correlation, which is visualized by **(I)** cartoon representation.

Based on interface cross-correlation result in Figure 5, sliced heat map was investigated in close view, followed by molecular dynamics simulation to understand how the effect of temperature on the dynamics and collectivity of certain critical residues involved in the allosteric regions (Fig. 6). Accordingly, effectively focused on the residues G1 and F221 (B chain, Phe1) then the residues V3 and N223 (B chain, Asn3). Residue F221 covers the allosteric regions of insulin B chain and has crucial role in allosteric shift the R-status to T-status; thus, directly affecting insulin activation as monomer (Maltesen et al., 2009). F221 (Phe1B) is also crucial for interaction with myristic acid of insulin detemir (Whittingham et al., 1997). Fluctuation of F221 through 50 ns is more stable and relatively stable at 100°K and 200°K dynamics, respectively, than that of ambient temperature. It aligns with cross-correlation matrices between G1-F221 residues that show high collectivity values close to 1 (red) in 100°K then 200°K dynamics. Ambient temperature (300°K) fluctuations of F221 became markedly mobile in simulations trend, which is closely correlated with reduced cross-correlation profile (G1-F221) that close to 0 (green). This shift suggests that elevated thermal motion disrupts the collective behavior of residues that critical for insulin allostery (Bonaccio et al., 2005; Huus et al., 2006). Likewise, the dynamics of V3 and N223 residues display similar trends. Especially, residue N223 (AsnB3) is highly significant to the scaffolding of hexamer stabilization through dimer-dimer electrostatic interactions (Palmieri et al., 2013), which is supported by our previous study showing alternate monomer kinetics of insulin due to disrupting AsnB3 interactions in the core of hexamer (Ayan et al., 2024). Cryogenic (100°K) and lower (200°K) temperatures show diminished fluctuations of N223 residue and high cross-correlations between V3-N223 residues, stressing AsnB3 role in supporting hexamer stability at those temperatures (Palmieri et al., 2013). The fluctuations of N223 residue at ambient temperature are higher by at least 15 Å through simulation, supported by loss of coherent movements for the V3-N223 coupling in cross-correlation. These findings suggest elevated temperatures might led to loosely coupled arrangement of hexameric structure, probably due to initializing the loss of noncovalent dimer-dimer (asu) relations (Espallargas et al., 2008; Song et al., 2013).

**Figure 6.**
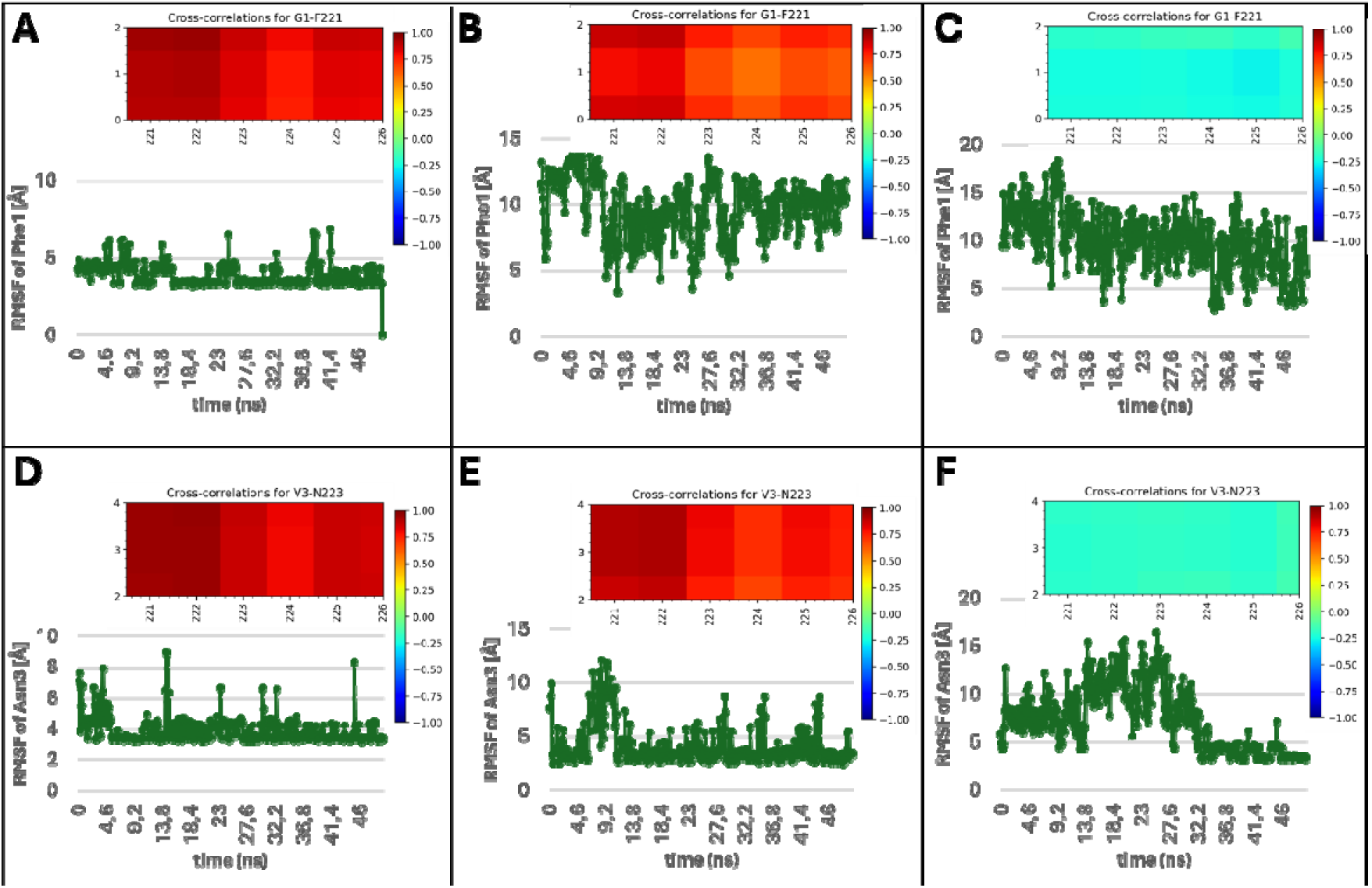
Molecular dynamics simulation along allosteric residues at interface region supports cross-correlation outputs. (**A, B, C**) F221 (Phe1 in B chain) fluctuations through 50 ns and collective motion from GNM between G1 in A chain and F221 residues. **(D, E, F)** N223 (Asn3 in B chain) fluctuations through 50 ns and collective motion from GNM between V3 in A chain and N223 residues. Toward elevated temperature, mobility and cooperativity change progressively. Y-axes of cross-correlation maps in panels A to C cover G1-I2 residues in chain A but focus solely on G1-F221 relations for discussion. Also, panels D to F cover I2-E4 residues in cross-correlation (y-axes) in chain A but focus solely on V3-N223 relations for discussion. RMSF was calculated for all atoms that builds Phe1 and Asn3 residues except hydrogens.

## Conclusion

We determined three crystal structures of dihexamer insulin (2.3 Å, 2.85 Å, and 2.88 Å resolutions) at 100°K, 200°K, and 300°K temperatures, respectively. For the first time, we observed a structural transition from the dihexamer to hexamer form at 200°K, with significant changes in the unit-cell parameters. The multi-temperature X-ray crystallographic structures were analyzed into two key aspects. **(i)** focused on the effects of temperature on the average protein structure and the changes in protein packing within the crystal lattice. **(ii)** investigated the influence of temperature on protein dynamics through experimental B-factor analysis, and computational NMA with MD simulations. Our findings reveal that sharp changes in temperature impact both the crystal lattice and mosaicity of insulin crystals. We hypothesize that the significant conformational shift from the dihexamer to hexamer structure is linked to the disruption of noncovalent interactions between the myristic acid moieties. We observed reduced collectivity and elevated thermal motion within the structure as the temperature increased. Residues critical for the allosteric transition of insulin exhibited apparent temperature-dependent alterations, as shown by MD simulations and NMA analyses. These findings provide valuable insights into insulin’s structural and dynamic behaviors across a broad temperature range, contributing to a deeper understanding of its thermal stability and allosteric mechanisms of a protracted insulin.

## Materials and Methods

### Sample Preparation and Crystallization

Detemir, marketed under the brand name Levemir®, was crystallized using the sitting-drop microbatch vapor diffusion screening technique under oil in 72-well Terasaki crystallization plates, as described by Ayan et al, (Ayan et al., 2023). Optimal crystals formed in a buffer containing 0.1 M MES monohydrate at pH 6.5, with 12% w/v polyethylene glycol 20,000 (Crystal Screen 2™, Hampton Research). The detemir solution was mixed in a 1:1 volume ratio (v/v) with this crystallization buffer at room temperature to initiate crystallization. Each well, containing 0.83 µL of protein and the crystallization buffer, was sealed with 16.6 µL of paraffin oil (Cat#ZS.100510.5000, ZAG Kimya, Turkey) and stored at 4°C until crystal harvesting. Crystal formation and growth were monitored under a compound light microscope, and large crystals developed within one week.

### Crystal harvesting

The optimal crystals were carefully collected from the Terasaki crystallization plates using MiTeGen and micro-loop sample pins attached to a magnetic wand under a compound light microscope. Once harvested, the crystals were promptly flash-frozen by immersing them in liquid nitrogen and then placed in a pre-cooled sample puck (Cat#M-CP-111-021, MiTeGen, Ithaca, NY, USA). The puck, kept in liquid nitrogen, was then transported to the dewar of the Turkish DeLight (Turkish Light Source, Turkey) for diffraction data collection.

### Data collection and processing

Diffraction data were acquired using Rigaku’s XtaLAB Synergy Flow XRD system equipped with CrysAlisPro software, following the protocol detailed by Atalay et al, (Atalay et al., 2022). Prior to data collection, the system was cooled to 100 K with an Oxford Cryosystems Cryostream 800 Plus. Liquid nitrogen was then added to the dewar, and a loaded sample puck was placed into the cryo sample storage dewar installed at the Turkish DeLight facility. The robotic auto-sample changer (UR3) utilized a six-axis robotic arm to mount the MiTeGen sample pin from the storage dewar onto the Intelligent Goniometer Head (IGH). The crystal was positioned at the X-ray interaction point using the CrysAlisPro software’s automatic centering feature (Figure S1). A 100 °K dataset was collected over two hours. Subsequently, the system temperature was adjusted to 200 °K with the Cryostream 800 Plus, and another two-hour dataset was collected. After completing the 100 °K and 200 °K datasets, the temperature was raised to 300 °K, with an extended overnight (5-hour) data collection due to the obtaining weaker diffraction patterns caused by the high temperature and radiation exposure.

The PhotonJet-R X-ray generator, equipped with a Cu X-ray source, operated at full power (40 kV and 30 mA), with beam intensity set to 10% via the piezo slit system to minimize cross-fire and prevent overlap of Bragg reflections. Diffraction data, totaling 33,832 reflections (100 °K), 48,296 reflections (200 °K), and 25,432 reflections (300 °K), were collected at a resolution of 2.3 Å, 2.85 Å, 2.88 Å, respectively, with a 45 mm detector distance, 0.2-degree scan width, and 1-second exposure time. The data were processed manually by the data reduction feature in CrysAlisPro. Once reduction was complete, the resulting files, containing both unmerged and unscaled data (.rrpprof format), were converted to structure factor files (.mtz format) via the *.hkl file. This conversion was facilitated by integrating the CCP4 crystallography suite with CrysAlisPro.

### Structure determination and refinement

The crystal structures of dihexamer and hexamer detemir were resolved to 2.3 Å, 2.85 Å, and 2.88 Å at 100 °K, 200 °K, and 300 °K, respectively, in space group R3: H. The unit cell dimensions at 100 °K were determined as a□=□b□=□c□=□79.06 Å, 533 α□=□β□=□90°, and γ□=□120° at a wavelength of 1.00 Å. At 200 °K and 300 °K, the unit cell dimensions shifted to a□=□b□=□81.24 Å, c□=□40.32 Å, α□=□β□=□90°, γ□=□120° and a□=□b□=□78.31 Å, c□=□39.63 Å, α□=□β□=□90°, γ□=_120°, respectively. Molecular replacement was conducted using the automated PHASER (McCoy et al., 2007) program in the PHENIX (Adams et al., 2010) software suite, with the previously determined cryogenic X-ray crystal structure (PDB ID: 1TRZ) serving as the reference model. The 1TRZ coordinates provided the basis for initial rigid-body refinement in PHENIX. Following simulated-annealing refinement, individual atomic coordinates and translation/libration/screw (TLS) parameters were further refined. Potential positions for altered side chains and water molecules were examined in COOT (Emsley et al., 2010), retaining those with significant difference densities. Missing water molecules were manually added, while those outside the electron density map were removed. Structure refinement statistics are presented in Table 1, and all figures for the X-ray crystal structures were created with PyMOL (Delano, 2002).

### Gaussian Network Model (GNM)

The Gaussian Network Model (GNM) analysis was conducted using ProDy (Bakan et al., 2011) on the detemir structures obtained 100 °K, 200 °K, and 300 °K temperatures. For each structure, contact maps were generated based on the Cα atoms, with a cutoff distance set at 8.00 Å. This yielded N − 1 nonzero normal modes, where N is the number of Cα atoms. Cross-correlations of residue fluctuations were calculated from the cumulative 10 slowest GNM modes, while squared fluctuations for each structure were derived from the weighted 10 fastest modes to assess local motions and dynamic behavior. Matplotlib was used to visualize normalized results graphically. Additionally, we performed network analysis of protein structures (NAPS) to examine node centrality, applying default settings on the NAPS Web server (Chakrabarty et al., 2016). B-factor normalization entails adjusting experimental B-factors to describe their distribution based on expected value and variance, facilitating comparison across different datasets by reducing dominant factors that could bias the data. During this normalization process, the pdb library stores the normalized values in a temporary record set, which contains B’-factors for each residue. The initial algorithm for B-factor normalization was introduced by Karplus and Schulz, linking the experimental B-factor of a specific residue to the arithmetic mean of all B-factors within a given structure: 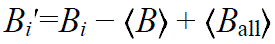

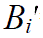: The normalized B-factor for a specific residue.

*B*_i_: The experimental B-factor of the specific residue.

⟨*B*⟩: The average B-factor of all residues within the structure.

*B*_all_: The overall average B-factor across all datasets used for normalization.

### Molecular dynamics (MD) simulation

Molecular dynamics (MD) simulations were conducted using the Nanoscale Molecular Dynamics (NAMD 3.0) (Phillips et al., 2005) program, employing the all-atom additive CHARMM36m force field (Huang et al., 2016) specifically designed for protein residues. MD was based on the work published by Anderson (Andersen, 1980). Simulations were performed under periodic boundary conditions to mimic an infinite system by allowing atoms exiting one side of the simulation box to re-enter from the opposite side, thus avoiding edge effects. In this setup, a time step of 2 femtoseconds (fs) was applied, and trajectory data was recorded at intervals of 10 picoseconds (ps). The initial temperature was set to 100 °K (cryogenic temperature) and maintained using a Langevin thermostat (Schneider et al., 1978) with parameters adjusted to keep the system stable (`langevinTemp`: 100 °K, `langevinDamping`: 1.0 ps ¹). Additionally, system pressure was regulated at 1.026 bar using a Langevin piston (`langevinPistonTarget`: 1.01325 atm) (Feller et al., 1995). For handling short-range interactions, electrostatic and van der Waals forces were cut off at 12 Å. To ensure that atoms moving close to the cutoff boundary remained within the interaction list without requiring frequent updates, the `pairlistdist` parameter was set to 14 Å, and a safety margin was established with a `skinnb` value of 2 Å. Sequential simulations at elevated temperatures were executed by preparing separate configuration files for each temperature stage (200 °K and 300 °K). The output from the 200 °K simulation served as the starting configuration for the 300 °K simulation. The system temperature was gradually raised from 100 °K to 200 °K and then to 300 °K over a period of 100 ps, using a harmonic restraint of 10.0 kcal/ (mol Å²) applied to the solution, thereby stabilizing the structural integrity during heating. Following the heating phase, a 5 nanosecond (ns) equilibration run was performed with an increased `skinnb` value to allow the system dimensions and density to stabilize. The temperature during equilibration and production runs was maintained using the Langevin thermostat under constant pressure periodic boundary conditions with an average pressure target of 1.01325 atm. Finally, 50 nanosecond production MD simulations were conducted with temperature control managed by the Langevin thermostat, ensuring a consistent thermodynamic environment throughout the simulation duration.

## Acknowledgments

The authors gratefully acknowledge the use of the services and facilities of the University of Health Sciences Experimental Medicine Research and Application Center (Validebag Research Park).

## Author contributions

E.A. and A.K. designed the experiment. E.A. wrote the entire main manuscript text performing all experiments, computational analyses and preparing all figures. E.A. and A.K. reviewed the manuscript.

## Data and materials availability

All data are available in the main text or the supplementary materials.

**Supplementary Figure 1.**
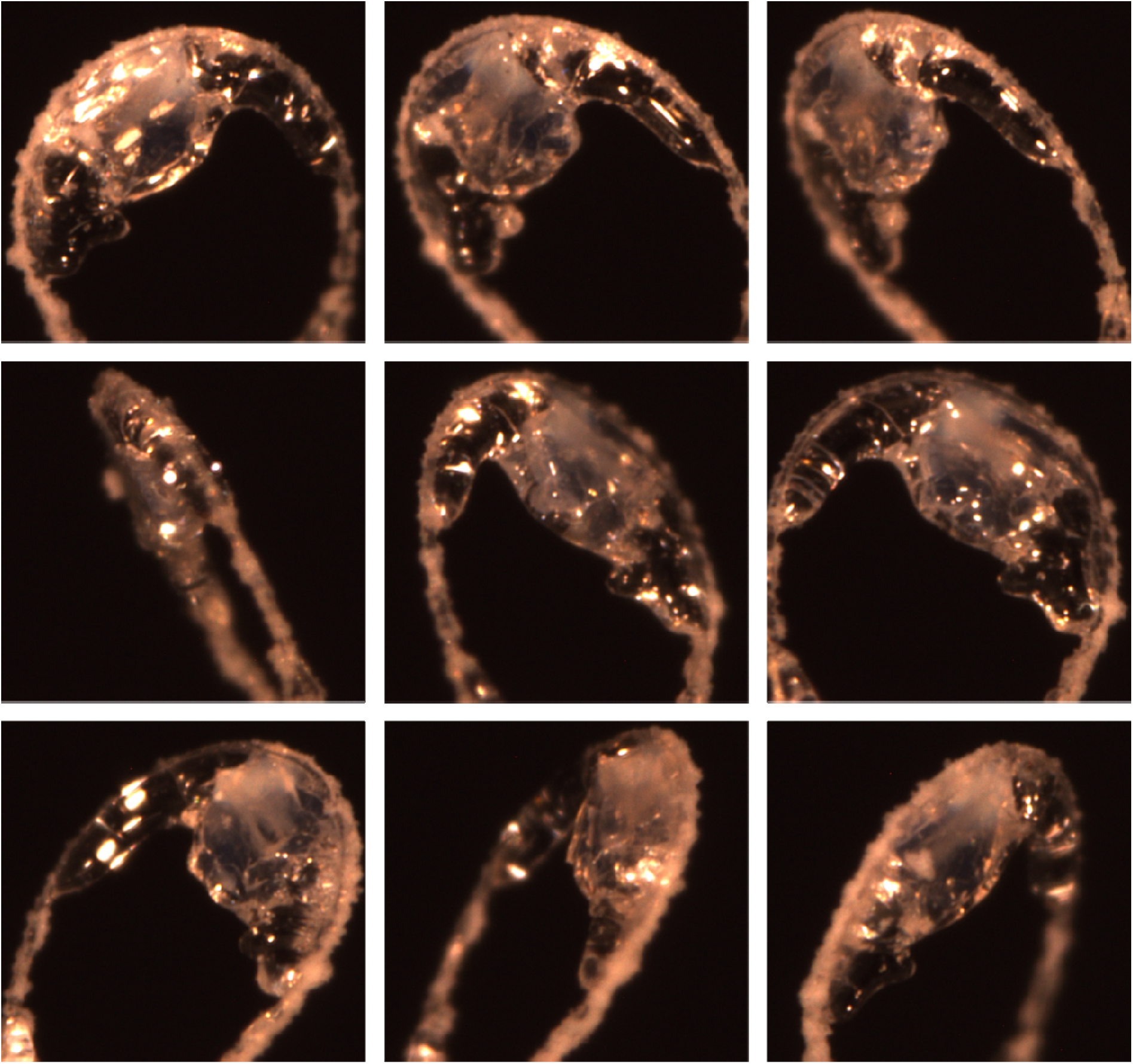
Single crystal on a loop for data collection through temperature-jump single crystal-X-ray crystallography. Representative images of the single insulin crystal used for X-ray data collection at 100 °K, 200 °K, and 300 °K temperatures. The crystal was maintained under cryogenic conditions for two hours at 100°K and 200 °K, followed by overnight data collection at 300 °K.

**Supplementary Figure 2.**
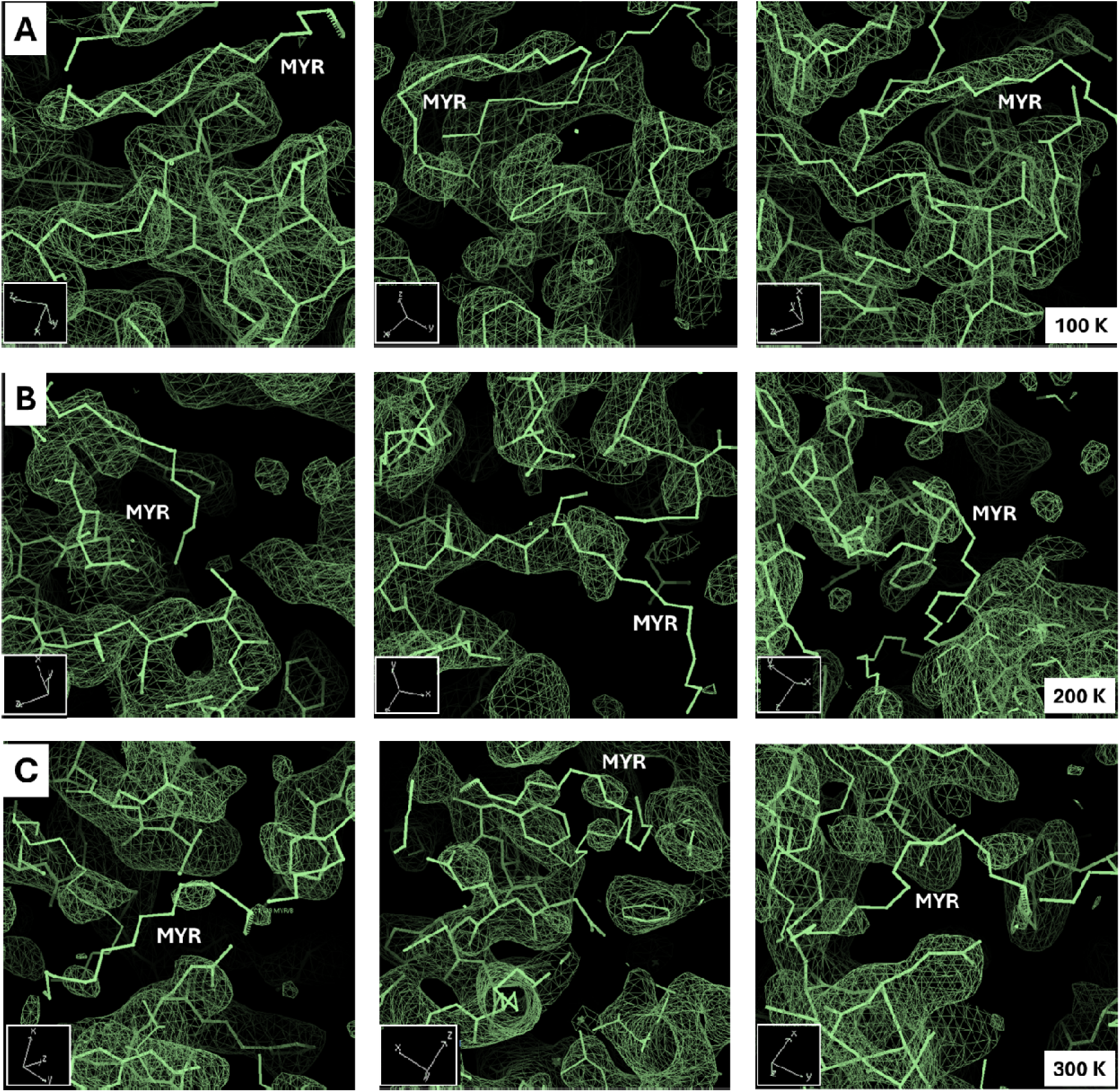
Temperature-dependent electron density maps of myristic acid in insulin crystals. Electron density maps (2*Fo-Fc*) of myristic acid (MYR) regions in insulin crystals at 100°K (A), 200°K (B), and 300°K (C). The maps are contoured at 1.0 σ and illustrate the structural integrity of MYR molecules across temperatures. At 100°K **(A)**, the electron density is well-defined, showing strong and continuous density along the MYR chains, indicative of their stable interactions with the surrounding residues. At 200°K **(B)**, partial disorder is observed, particularly at the distal ends of MYR, suggesting increased flexibility, correlating with the transition from the dihexameric to hexameric state. At 300°K **(C)**, the electron density becomes fragmented, reflecting significant displacement and dynamic behavior of MYR. Insets highlight the orientation of MYR molecules within their respective binding pockets.

## References

1. Adams, P. D., Afonine, P. V., Bunkóczi, G., Chen, V. B., Davis, I. W., Echols, N., Headd, J. J., Hung, L. W., Kapral, G. J., Grosse-Kunstleve, R. W., McCoy, A. J., Moriarty, N. W., Oeffner, R., Read, R. J., Richardson, D. C., Richardson, J. S., Terwilliger, T. C., & Zwart, P. H. (2010). PHENIX: A comprehensive Python-based system for macromolecular structure solution. Acta Crystallographica Section D: Biological Crystallography. doi: 10.1107/S0907444909052925

2. Andersen, H. C. (1980). Molecular dynamics simulations at constant pressure and/or temperature. The Journal of Chemical Physics. doi: 10.1063/1.439486

3. Atalay, N., Akcan, E. K., Gül, M., Ayan, E., Destan, E., Ertem, F. B., Tokay, N., Çakilkaya, B., Nergiz, Z., Karakadioğlu, G., Kepceoğlu, A., Yapici, İ., Tosun, B., Baldir, N., Yildirim, G., Johnson, J. A, Güven, Ö., Sğhafiei, A., Arslan, N. E., … Demirci, H. (2022). Cryogenic X-ray crystallographic studies of biomacromolecules at Turkish Light Source “Turkish DeLight.” BioRxiv, 10, 2022.09.03.506456. doi: 10.1101/2022.09.03.506456

4. Atilgan, A. R., Durell, S. R., Jernigan, R. L., Demirel, M. C., Keskin, O., & Bahar, I. (2001). Anisotropy of fluctuation dynamics of proteins with an elastic network model. Biophysical Journal. doi: 10.1016/S0006-3495(01)76033-X

5. Ayan, E., & DeMirci, H. (2022). A Brief Atlas of Insulin. Current Diabetes Reviews, 19. doi: 10.2174/1573399819666220610150342

6. Ayan, E., Destan, E., Kepceoglu, A., Ciftci, H. I., Kati, A., & DeMirci, H. (2023). Comparative Study of High-Resolution LysB29(Nε-myristoyl) des(B30) Insulin Structures Display Novel Dynamic Causal Interrelations in Monomeric-Dimeric Motions. Crystals. doi: 10.3390/cryst13040648

7. Ayan, E., Türk, M., Tatlı, Ö., Bostan, S., Telek, E., Dingiloğlu, B., Dogan, B. Z., Alp, M. I., Katı, A., Dinler-Doğanay, G., & DeMirci, H. (2024). X-ray Crystallographic and Hydrogen Deuterium Exchange Studies Confirm Alternate Kinetic Models for Homolog Insulin Monomers. BioRxiv, 2024.04.16.589767. doi: 10.1101/2024.04.16.589767

8. Ayan, E., Yuksel, B., Destan, E., Ertem, F. B., Yildirim, G., Eren, M., Yefanov, O. M., Barty, A., Tolstikova, A., Ketawala, G. K., Botha, S., Dao, E. H., Hayes, B., Liang, M., Seaberg, M. H., Hunter, M. S., Batyuk, A., Mariani, V., Su, Z., … DeMirci, H. (2022). Cooperative allostery and structural dynamics of streptavidin at cryogenic– and ambient-temperature. Communications Biology. doi: 10.1038/s42003-021-02903-7

9. Bagler, G., & Sinha, S. (2007). Assortative mixing in protein contact networks and protein folding kinetics. Bioinformatics. doi: 10.1093/bioinformatics/btm257

10. Bahar, I., Atilgan, A. R., Demirel, M. C., & Erman, B. (1998). Vibrational dynamics of folded proteins: Significance of slow and fast motions in relation to function and stability. Physical Review Letters. doi: 10.1103/PhysRevLett.80.2733

11. Bahar, I., Atilgan, A. R., & Erman, B. (1997). Direct evaluation of thermal fluctuations in proteins using a single-parameter harmonic potential. Folding and Design. doi: 10.1016/S1359-0278(97)00024-2

12. Baheri, M., & Dayer, M. R. (2016). Temperature and pH Effects on Insulin Structure: A Molecular Dynamic Approach. Jentashapir Journal of Health Research. doi: 10.17795/jjhr-36931

13. Bakan, A., Meireles, L. M., & Bahar, I. (2011). ProDy: Protein dynamics inferred from theory and experiments. Bioinformatics. doi: 10.1093/bioinformatics/btr168

14. Bech, E. M., Pedersen, S. L., & Jensen, K. J. (2018). Chemical Strategies for Half-Life Extension of Biopharmaceuticals: Lipidation and Its Alternatives. In ACS Medicinal Chemistry Letters. doi: 10.1021/acsmedchemlett.8b00226

15. Bedford, A., Beckett, L., Harthan, L., Wang, C., Jiang, N., Schramm, H., Guan, L. L., Daniels, K. M., Hanigan, M. D., & White, R. R. (123 C.E.). Ruminal volatile fatty acid absorption is affected by elevated ambient temperature. doi: 10.1038/s41598-020-69915-x

16. Blacklock, K., & Verkhivker, G. M. (2014). Computational Modeling of Allosteric Regulation in the Hsp90 Chaperones: A Statistical Ensemble Analysis of Protein Structure Networks and Allosteric Communications. PLoS Computational Biology. doi: 10.1371/journal.pcbi.1003679

17. Bonaccio, M., Ghaderi, N., Borchardt, D., & Dunn, M. F. (2005). Insulin allosteric behavior: Detection, identification, and quantification of allosteric states via 19F NMR. Biochemistry. doi: 10.1021/bi050392s

18. Brange, J., & Langkjoer, L. (1993). Insulin structure and stability. In Pharmaceutical biotechnology (Vol. 5, pp. 315–350). Springer, Boston, MA. doi: 10.1007/978-1-4899-1236-7_11

19. Brinda, K. V., & Vishveshwara, S. (2005). A network representation of protein structures: Implications for protein stability. Biophysical Journal. doi: 10.1529/biophysj.105.064485

20. Chakrabarty, B., & Parekh, N. (2016). NAPS: Network analysis of protein structures. Nucleic Acids Research. doi: 10.1093/nar/gkw383

21. Ciszak, E., & Smith, G. D. (1994). Crystallographic Evidence for Dual Coordination around Zinc in the T3R3 Human Insulin Hexamer. Biochemistry. doi: 10.1021/bi00172a030

22. Cucka, P., Singman, L., Lovell, F. M., & Low, B. W. (1970). Studies of insulin crystals at low temperatures. Effects on lattice dimensions, temperature parameters and structure. Acta Crystallographica Section B Structural Crystallography and Crystal Chemistry. doi: 10.1107/s0567740870004843

23. Delano, W. L. (2002). The PyMOL Molecular Graphics System. CCP4 Newsletter on Protein Crystallography.

24. Demirel, M. C., Atilgan, A. R., Jernigan, R. L., Erman, B., & Bahar, I. (1998). Identification of kinetically hot residues in proteins. Protein Science. doi: 10.1002/pro.5560071205

25. Emsley, P., Lohkamp, B., Scott, W. G., & Cowtan, K. (2010). Features and development of Coot. Acta Crystallographica Section D: Biological Crystallography. doi: 10.1107/S0907444910007493

26. Eren, M., Tuncbag, N., Jang, H., Nussinov, R., Gursoy, A., & Keskin, O. (2021). Normal Mode Analysis of KRas4B Reveals Partner Specific Dynamics. Journal of Physical Chemistry B. doi: 10.1021/acs.jpcb.1c00891

27. Espallargas, G. M., Brammer, L., Allan, D. R., Pulham, C. R., Robertson, N., & Warren, J. E. (2008). Noncovalent interactions under extreme conditions: High-pressure and low-temperature diffraction studies of the isostructural metal-organic networks (4-chloropyridinium)2[CoX4] (X = Cl, Br). Journal of the American Chemical Society. doi: 10.1021/ja8010868

28. Feller, S. E., Zhang, Y., Pastor, R. W., & Brooks, B. R. (1995). Constant pressure molecular dynamics simulation: The Langevin piston method. The Journal of Chemical Physics. doi: 10.1063/1.470648

29. Fischer, M. (2021). Macromolecular room temperature crystallography. In Quarterly Reviews of Biophysics. doi: 10.1017/S0033583520000128

30. Frauenfelder, H., Hartmann, H., Parak, F., Karplus, M., Kuriyan, J., Kuntz, I. D., Tilton, R. F., Petsko, G. A., Ringe, D., Connolly, M. L., & Max, N. (1987). Thermal Expansion of a Protein. Biochemistry. doi: 10.1021/bi00375a035

31. Frauenfelder, H., Petsko, G. A., & Tsernoglou, D. (1979). Temperature-dependent x-ray diffraction as a probe of protein structural dynamics. Nature. doi: 10.1038/280558a0

32. Gursky, O., Badger, J., Li, Y., & Caspar, D. L. (1992). Conformational changes in cubic insulin crystals in the pH range 7–11. Biophysical Journal. doi: 10.1016/S0006-3495(92)81697-1

33. Hartmann, H., Parak, F., Steigemann, W., Petsko, G. A., Ponzi, D. R., & Frauenfelder, H. (1982). Conformational substates in a protein: structure and dynamics of metmyoglobin at 80 K. Proceedings of the National Academy of Sciences of the United States of America. doi: 10.1073/pnas.79.16.4967

34. Hassiepen, U., Federwisch, M., Mülders, T., & Wollmer, A. (1999). The lifetime of insulin hexamers. Biophysical Journal. doi: 10.1016/S0006-3495(99)77012-8

35. Holleman, F. (2015). Insulin lispro (revision number 20). In Diapedia. doi: 10.14496/dia.8104096170.20

36. Home, P., & Kurtzhals, P. (2006). Insulin detemir: From concept to clinical experience. In Expert Opinion on Pharmacotherapy. doi: 10.1517/14656566.7.3.325

37. Huang, J., Rauscher, S., Nawrocki, G., Ran, T., Feig, M., De Groot, B. L., Grubmüller, H., & MacKerell, A. D. (2016). CHARMM36m: An improved force field for folded and intrinsically disordered proteins. Nature Methods. doi: 10.1038/nmeth.4067

38. Huus, K., Havelund, S., Olsen, H. B., Sigurskjold, B. W., Van De Weert, M., & Frokjaer, S. (2006). Ligand binding and thermostability of different allosteric states of the insulin zinc-hexamer. Biochemistry. doi: 10.1021/bi0524520

39. Jarosinski, M. A., Dhayalan, B., Chen, Y. S., Chatterjee, D., Varas, N., & Weiss, M. A. (2021). Structural principles of insulin formulation and analog design: A century of innovation. In Molecular Metabolism. doi: 10.1016/j.molmet.2021.101325

40. Juers, D. H., & Matthews, B. W. (2001). Reversible lattice repacking illustrates the temperature dependence of macromolecular interactions. Journal of Molecular Biology. doi: 10.1006/jmbi.2001.4891

41. Kihara, K. (1978). Thermal change in unit-cell dimensions, and a hexagonal structure of tridymite. Zeitschrift Fur Kristallographie – New Crystal Structures. doi: 10.1524/zkri.1978.148.3-4.237

42. Kontopoulos, D.-G., Patmanidis, I., Barraclough, T. G., & Pawar, S. (2020). Higher temperatures worsen the effects of mutations on protein stability. BioRxiv, 2020.10.13.337972. doi: 10.1101/2020.10.13.337972

43. Kuczera, K., Kuriyan, J., & Karplus, M. (1990). Temperature dependence of the structure and dynamics of myoglobin. A simulation approach. Journal of Molecular Biology. doi: 10.1016/S0022-2836(05)80196-2

44. Kurinov, I. V., & Harrison, R. W. (1995). The influence of temperature on lysozyme crystals. Structure and dynamics of protein and water. Acta Crystallographica – Section D Biological Crystallography. doi: 10.1107/S0907444994009261

45. Kuzmanic, A., & Zagrovic, B. (2010). Determination of ensemble-average pairwise root mean-square deviation from experimental B-factors. Biophysical Journal. doi: 10.1016/j.bpj.2009.11.011

46. Lakhloufi, S., Tailleur, E., Guo, W., Le Gac, F., Marchivie, M., Lemée-Cailleau, M. H., Chastanet, G., & Guionneau, P. (2018). Mosaicity of Spin-crossover crystals. Crystals. doi: 10.3390/cryst8090363

47. Maltesen, M. J., Bjerregaard, S., Hovgaard, L., Havelund, S., & Van De Weert, M. (2009). Analysis of insulin allostery in solution and solid state with FTIR. Journal of Pharmaceutical Sciences. doi: 10.1002/jps.21736

48. Massana-Cid, H., Maggi, C., Gnan, N., Frangipane, G., & Di Leonardo, R. (2024). Multiple temperatures and melting of a colloidal active crystal. Nature Communications 2024 15:1, 15(1), 1–9. doi: 10.1038/s41467-024-50937-2

49. McCoy, A. J., Grosse-Kunstleve, R. W., Adams, P. D., Winn, M. D., Storoni, L. C., & Read, R. J. (2007). Phaser crystallographic software. Journal of Applied Crystallography. doi: 10.1107/S0021889807021206

50. Medeiros Almeida, V., Chaudhuri, A., Cangussu Cardoso, M. V., Matsuyama, B. Y., Monteiro Ferreira, G., Goulart Trossini, G. H., Salinas, R. K., Loria, J. P., & Marana, S. R. (2021). Role of a high centrality residue in protein dynamics and thermal stability. Journal of Structural Biology. doi: 10.1016/j.jsb.2021.107773

51. Meents, A., Gutmann, S., Wagner, A., & Schulze-Briese, C. (2010). Origin and temperature dependence of radiation damage in biological samples at cryogenic temperatures. Proceedings of the National Academy of Sciences of the United States of America. doi: 10.1073/pnas.0905481107

52. Novikov, V. V., Matovnikov, A. V., Mitroshenkov, N. V., Shevelkov, A. V., & Bud’Ko, S. L. (2020). Crystal lattice disorder and characteristic features of the low-temperature thermal properties of higher borides. Dalton Transactions. doi: 10.1039/c9dt04919c

53. Palmieri, L. C., Fávero-Retto, M. P., Lourenço, D., & Lima, L. M. T. R. (2013). A T3R3 hexamer of the human insulin variant B28Asp. Biophysical Chemistry. doi: 10.1016/j.bpc.2013.01.003

54. Parak, F., Hartmann, H., Aumann, K. D., Reuscher, H., Rennekamp, G., Bartunik, H., & Steigemann, W. (1987). Low temperature X-ray investigation of structural distributions in myoglobin. European Biophysics Journal. doi: 10.1007/BF00577072

55. Parak, F., Knapp, E. W., & Kucheida, D. (1982). Protein dynamics. Mössbauer spectroscopy on deoxymyoglobin crystals. Journal of Molecular Biology. doi: 10.1016/0022-2836(82)90285-6

56. Phillips, J. C., Braun, R., Wang, W., Gumbart, J., Tajkhorshid, E., Villa, E., Chipot, C., Skeel, R. D., Kalé, L., & Schulten, K. (2005). Scalable molecular dynamics with NAMD. In Journal of Computational Chemistry. doi: 10.1002/jcc.20289

57. Rasmussen, B. F., Stock, A. M., Ringe, D., & Petsko, G. A. (1992). Crystalline ribonuclease a loses function below the dynamical transition at 220 K. Nature. doi: 10.1038/357423a0

58. Rasmussen, M., Hach, M., Iwersen, J., Anil, G., Rosborg, H., Larsen, J., Hoffmann, L. C., & Kurtzhals, P. (2024). The Human Insulin Thermal Solution project—a private sector initiative to address the thermostability of insulin. The Lancet Diabetes and Endocrinology, 12(5), 292–294. doi: 10.1016/S2213-8587(24)00094-9

59. Rosskamp, R. H., & Park, G. (1999). Long-acting insulin analogs. Diabetes Care.

60. Saldivar-Garcia, A. J., & Lopez, H. F. (2004). Temperature effects on the lattice constants and crystal structure of a Co-27Cr-5Mo low-carbon alloy. Metallurgical and Materials Transactions A: Physical Metallurgy and Materials Science. doi: 10.1007/s11661-006-0232-6

61. Samuel, D., Ganesh, G., Yang, P.-W., Chang, M.-M., Wang, S.-L., Hwang, K.-C., Yu, C., Jayaraman, G., Kumar, T. K. S., Trivedi, V. D., & Chang, D.-K. (2008). Proline inhibits aggregation during protein refolding. Protein Science. doi: 10.1110/ps.9.2.344

62. Sarı, A., & Kaygusuz, K. (2001). Thermal performance of myristic acid as a phase change material for energy storage application. Renewable Energy. doi: 10.1016/S0960-1481(00)00167-1

63. Schneider, T., & Stoll, E. (1978). Molecular-dynamics study of a three-dimensional one-component model for distortive phase transitions. Physical Review B. doi: 10.1103/PhysRevB.17.1302

64. Smith, G. D., Ciszak, E., Magrum, L. A., Pangborn, W. A., & Blessing, R. H. (2000). R6 hexameric insulin complexed with m-cresol or resorcinol. Acta Crystallographica Section D: Biological Crystallography. doi: 10.1107/S0907444900012749

65. Somero, G. N. (2003). Protein adaptations to temperature and pressure: Complementary roles of adaptive changes in amino acid sequence and internal milieu. Comparative Biochemistry and Physiology – B Biochemistry and Molecular Biology. doi: 10.1016/S1096-4959(03)00215-X

66. Song, S., Dong, R., Wang, D., Song, A., & Hao, J. (2013). Temperature regulated supramolecular structures via modifying the balance of multiple non-covalent interactions. Soft Matter. doi: 10.1039/c3sm00006k

67. Stevens, L. A., Goetz, K. P., Fonari, A., Shu, Y., Williamson, R. M., Brédas, J. L., Coropceanu, V., Jurchescu, O. D., & Collis, G. E. (2015). Temperature-mediated polymorphism in molecular crystals: The impact on crystal packing and charge transport. Chemistry of Materials. doi: 10.1021/cm503439r

68. Sun, Q., Fu, Y., & Wang, W. (2022). Temperature effects on hydrophobic interactions: Implications for protein unfolding. Chemical Physics. doi: 10.1016/j.chemphys.2022.111550

69. Taylor, N. R. (2013). Small world network strategies for studying protein structures and binding. In Computational and Structural Biotechnology Journal. doi: 10.5936/csbj.201302006

70. Thermostability of human insulin Background (n.d.).

71. Tilton, R. F., Dewan, J. C., & Petsko, G. A. (1992). Effects of Temperature on Protein Structure and Dynamics: X-ray Crystallographic Studies of the Protein Ribonuclease-A at Nine Different Temperatures from 98 to 320 K. Biochemistry, 31(9), 2469–2481. doi: 10.1021/BI00124A006/ASSET/BI00124A006.FP.PNG_V03

72. Vimalavathini, R., & Gitanjali, B. (2009). Effect of temperature on the potency & pharmacological action of insulin. Indian Journal of Medical Research.

73. Wang, Z., Li, Y., Jiang, L., Qi, B., & Zhou, L. (2014). Relationship between secondary structure and surface hydrophobicity of soybean protein isolate subjected to heat treatment. Journal of Chemistry. doi: 10.1155/2014/475389

74. Whittingham, J. L., Havelund, S., & Jonassen, I. (1997). Crystal structure of a prolonged-acting insulin with albumin-binding properties. Biochemistry. doi: 10.1021/bi9625105

75. Yang, L. W., & Bahar, I. (2005). Coupling between catalytic site and collective dynamics: A requirement for mechanochemical activity of enzymes. Structure. doi: 10.1016/j.str.2005.03.015

76. Young, A. C. M., Tilton, R. F., & Dewan, J. C. (1994). Thermal expansion of hen egg-white lysozyme. Comparison of the 1·9 Å resolution structures of the tetragonal form of the enzyme at 100 K and 298 K. Journal of Molecular Biology. doi: 10.1016/S0022-2836(05)80034-8

77. Zheng, J., Guo, N., Huang, Y., Guo, X., & Wagner, A. (2024). High temperature delays and low temperature accelerates evolution of a new protein phenotype. Nature Communications 2024 15:1, 15(1), 1–14. doi: 10.1038/s41467-024-46332-6

78. Zhuravleva, M., Lindsey, A., Chakoumakos, B. C., Custelcean, R., Meilleur, F., Hughes, R. W., Kriven, W. M., & Melcher, C. L. (2015). Crystal structure and thermal expansion of a CsCe2Cl7 scintillator. Journal of Solid State Chemistry, 227, 142–149. doi: 10.1016/J.JSSC.2015.03.032.

